# Development and Application of a High-Content Virion Display Human GPCR Array

**DOI:** 10.1101/377754

**Authors:** Guan-Da Syu, Shih-Chin Wang, Guangzhong Ma, Shuang Liu, Donna Pearce, Atish Prakash, Brandon Henson, Lien-Chun Weng, Devlina Ghosh, Pedro Ramos, Daniel Eichinger, Ignacio Pino, Xinzhong Dong, Jie Xiao, Shaopeng Wang, Nongjian Tao, Kwang Sik Kim, Prashant J. Desai, Heng Zhu

**Affiliations:** Department of Pharmacology and Molecular Sciences, Johns Hopkins University School of Medicine, Baltimore, MD 21205, U.S.A.; Center for High-Throughput Biology, Johns Hopkins University School of Medicine, Baltimore, MD 21205, U.S.A.; Viral Oncology Program, Department of Oncology, The Sidney Kimmel Comprehensive Cancer Center, Johns Hopkins University School of Medicine, Baltimore, MD 21231, U.S.A.; Department of Biophysics and Biophysical Chemistry, Johns Hopkins University School of Medicine, Baltimore, MD 21205, U.S.A.; Biodesign Center for Bioelectronics and Biosensors, Arizona State University, Tempe, AZ 85287, U.S.A.; Division of Paediatric Infectious Diseases, Johns Hopkins University School of Medicine, Baltimore, MD 21287, U.S.A.; CDI Laboratories, Inc. Mayaguez, Puerto Rico 00682, U.S.A.; The Solomon H. Snyder Department of Neuroscience, Center for Sensory Biology, Johns Hopkins University School of Medicine, Baltimore, MD 21205, U.S.A.; Howard Hughes Medical Institute, Johns Hopkins University School of Medicine, Baltimore, MD 21205, U.S.A.; School of Electrical, Computer and Energy Engineering, Arizona State University, Tempe, AZ 85287, U.S.A.

## Abstract

G protein-coupled receptors (GPCRs) comprise the largest membrane protein family in humans and can respond to a wide variety of ligands and stimuli. Like other multi-pass membrane proteins, the biochemical properties of GPCRs are notoriously difficult to study because they must be embedded in lipid bilayers to maintain their native conformation and function. To enable an unbiased, high-throughput platform to profile biochemical activities of GPCRs in native conformation, we individually displayed 315 human non-odorant GPCRs (>85% coverage) in the envelope of human herpes simplex virus-1 and immobilized on glass to form a high-content <u>Vir</u>ion <u>D</u>isplay (VirD) array. Using this array, we found that 50% of the tested commercial anti-GPCR antibodies (mAbs) is ultra-specific, and that the vast majority of those VirD-GPCRs, which failed to be recognized by the commercial mAbs, could bind to their canonical ligands, indicating that they were folded correctly. Next, we used the VirD-GPCR arrays to examine binding specificity of two known peptide ligands and recovered expected interactions, as well as new off-target interactions, three of which were confirmed with real-time kinetics measurements. Finally, we explored the possibility of discovering novel pathogen targets by probing VirD-GPCR arrays with live group B Streptococcus (GBS), a common Gram-positive bacterium causing neonatal meningitis. Using cell invasion assays and a mouse model of hematogenous meningitis, we showed that inhibition of one of the five newly identified GPCRs, CysLTR1, greatly reduced GBS penetration in brain-derived endothelial cells and in mouse brains. Therefore, our work demonstrated that the VirD-GPCR array holds great potential for high-throughput, unbiased screening for small molecule drugs, affinity reagents, and deorphanization.

## Introduction

The G protein-coupled receptor (GPCR) family is comprised of >800 members, forming the largest membrane protein family in humans. Almost all of the annotated GPCRs function via binding to agonists or antagonists, ranging from ions to metabolites, small molecules, peptides and proteins, and exhibit a wide variety of signaling pathways. As archived in the IUPHAR/BPS database, GPCRs are classified into three large classes (i.e., Classes A, B, and C) and two smaller classes (i.e., Frizzled and Adhesion). Of all annotated human GPCRs, 370 are non-odorant and play crucial roles in many different aspects of cellular processes. They are the most preferred drug targets because their dysregulation often lead to disease or cancer. In fact, of the ~1,400 FDA-approved drugs, 475 (34%) target 108 non-odorant GPCRs ^1–3^. Although 70% of them have known ligands, a significant fraction of them are “orphans” without an identified ligand ^1–3^.

However, non-odorant GPCRs are notoriously difficult to study for several reasons. First, GPCRs must be embedded in a membrane to maintain proper conformation, and many also require a series of posttranslational modifications, such as glycosylation, for activity ^4^. Second, despite the recent progress in the cloning and expression of GPCR proteins in cells, their expression levels vary greatly, hampering an effective strategy to study these proteins in parallel and on a genomic scale ^5^. Third, a considerable number of GPCRs are toxic to the host cells once overexpressed, making it difficult to establish stable cell lines that constitutively express these receptors ^6^. Finally and perhaps most importantly, 116 (31%) of the ~370 non-odorant GPCRs are orphans, without known physiological agonists or antagonists, which precludes study of their activity.

Because of the significant importance of these “druggable” GPCRs, it is critical to develop scalable technology platforms to enable their characterizations in a high-throughput fashion. Human protein arrays carrying GPCRs are available from several commercial sources, such as Invitrogen (Life Sciences), Origen Inc., and CDI Laboratories. However, GPCRs spotted on these commercial arrays are unlikely to maintain their native conformation or activities because detergents were used to purify these proteins. In a series of publications, Fang, Burkhalter and colleagues at Corning Inc. described fabrication of human GPCR arrays by spotting down membrane preps of mammalian cells in which a particular GPCR-of-interest was overexpressed ^7, 8^. However, membrane preparations used to fabricate the GPCR arrays are unavoidably contaminated with other endogenously expressed GPCRs, raising a homogeneity issue. Moreover, it is almost impossible to control concentration or orientation of the GPCRs on the array surface. Finally, because of stability issues caused by spotting fractured membrane preps, oligosaccharides were often added to associate with the head groups of the lipid bilayers, resulting in a higher susceptibility to buffer composition and pH.

As a proof-of-principle, we developed Virion Display (VirD) technology with which both single (i.e., CD4) and multi-pass (i.e., GPR77) membrane proteins could be displayed on the envelopes of herpes simplex virus-1 (HSV-1) with correct orientation and conformation ^9^. To demonstrate the power of VirD technology in this study, we fabricated a high-content VirD-GPCR array, comprised of 315 non-odorant GPCRs, and demonstrated that the VirD-GPCRs were folded and functional using a variety of biochemical assays.

## RESULTS

### Fabrication of a non-odorant VirD-GPCR array

To develop the first high-content VirD array, we decided to focus on the 370 non-odorant GPCRs. We assembled a collection of open reading frames (ORFs) encoding 337 non-odorant GPCRs that were available to us. To enable high-throughput cloning of these ORFs into the HSV-1 strain KOS genome carried in a bacterial artificial chromosome (BAC) vector ^10^, we replaced the glycoprotein B coding sequence (*UL27*) with a Gateway cloning cassette. The engineered molecule also contained the sequences for the expression of a V5 epitope at the C-terminus of the cloned ORF followed by a STOP codon (**Fig. 1a**). Meanwhile, STOP codons of the GPCR ORFs were removed via PCR reactions and the modified ORFs subcloned into the Gateway Entry vector. Single colonies were picked, followed by Sanger sequencing to ensure error-free subcloning of each ORF. Confirmed STOP-free ORFs were then shuttled into the *UL27* (gB) locus in the HSV-1 genome using LR recombination reactions and transformed into *E. coli* by electroporation. For each bacterial transformation, at least two colony PCR reactions were performed using a primer pair that annealed to the viral sequences flanking the cloning site of the GPCR ORFs, and gel electrophoresis was employed to examine whether the amplicon was of the expected size of the GPCR cloned. To this end, we have successfully subcloned a total of 332 (98.5%) GPCR ORFs into the *UL27* locus.

**Figure 1.**
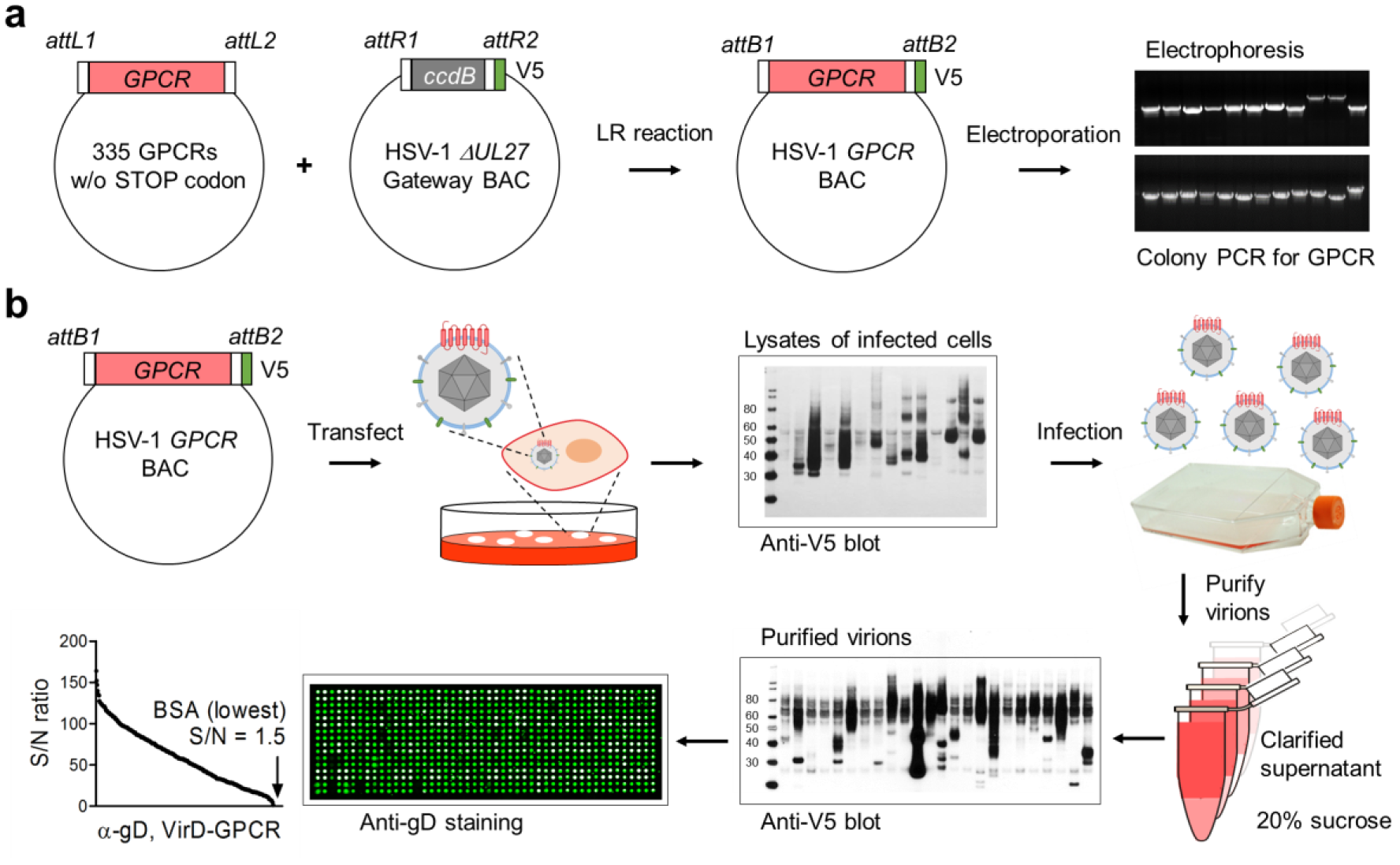
Construction of high-content VirD-GPCR array. (**a**) Subcloning of 335 human GPCR ORFs into the *UL27* locus of the HSV-1 genome. After the STOP codons were removed from the 335 available GPCR ORFs, they were subcloned into the *UL27* locus in the HSV-1 genome on a BAC vector, resulting in fusion with a V5 tag at their C-termini (middle panel). After bacterial transformation, colony PCR reactions were carried out and the products examined using electrophoresis to identify the correct construct (right panel). (**b**) Production of VirD-GPCR virions and VirD array fabrication. Confirmed recombinant virus constructs were individually transfected to Vero cells and the viruses were harvested ~7 days post-transfection. Anti-V5 mAb was used to examine expression of the GPCRs as a quality control. Passers were next used to infect cells for virion production. After sucrose cushion centrifugation, a fraction of purified virions were examined again with anti-V5. 315 VriD-GPCRs passed this quality control step and were spotted onto a glass slide to form VirD-GPCR array. The quality of VirD-GPCR array was examined using anti-gD mAb, followed by a Cy3-labeled anti-mouse IgG antibody. All the ViP.rD-GPCRs on the array showed positive anti-gD signals while the BSA showed the lowest signals.

To produce recombinant viruses, we first transfected each *GPCR:: ΔUL27* BAC DNA into a Vero transformed cell line, D87, that complements the growth of mutants that do not express gB. When viral plaques became evident, 5-7 days post-transfection, low titer viral stocks were harvested for each GPCR recombinant virus. After a secondary infection, expression of a total of 317 GPCRs was detected in total cell lysates with anti-V5 antibodies (**Fig. 1b**). Next, high titer stocks of the 317 VirD-GPCRs were individually prepared following infection of D87 cells. Since we observed that expression levels of different GPCRs varied in different cell lines, we prepared the final VirD-GPCR virions from Vero-, HEL-, HeLa-, and HEK293T-infected cells to maximize the production of VirD-GPCRs (**Supplementary Fig. 1a-b**). The VirD-GPCR virions were further purified to homogeneity via sucrose cushion and resuspended in a small volume to maintain a high virion concentration. A small fraction of each purified VirD-GPCR virion was subjected to anti-V5 immunoblot analysis, based on which 315 VirD-GPCRs passed this quality control step (**Fig. 1b**). Finally, the 315 virion preparations were arrayed into a 384-well titer dish and robotically printed in duplicate onto SuperEpoxy slides to form the VirD-GPCR array. The quality of the printed VirD-GPCR arrays was examined with anti-gD antibody and all of the 315 arrayed VirD-GPCRs showed significant anti-gD signals as compared with the negative controls (e.g., BSA) (lower left panel; **Fig. 1b**). Moreover, scatter plot analysis of an anti-gD assay performed in duplicate indicated a high reproducibility with a correlation coefficient of 0.92 (**Supplementary Fig. 1c-d**). Therefore, we successfully produced a high-content VirD array that covers 85% of the annotated non-odorant human GPCRs.

### Profiling antibody specificity on VirD-GPCR array

Antibody-based biologicals are emerging as the next generation therapeutics because of their unique properties, such as superior pharmacokinetics, simple formulation, and modular, easily engineerable format. Indeed, Erenumab (trade name Aimovig) was recently approved by the FDA as the first antibody-based drug that targets the GPCR, i.e., CGRPR, for the prevention of migraine ^11^. However, there are serious problems with the quality and consistency of antibodies because of the absence of standardized antibody-validation criteria, a lack of transparency from commercial antibody suppliers and technical difficulties in comprehensive assessments of antibody cross-reactivity ^12–17^.

To demonstrate that the VirD-GPCR arrays can be used to test antibody specificity, we selected 20 commercially available mAbs targeting 20 human GPCRs. Since all of these mAbs are documented by the commercial providers to recognize intact cells (e.g., flow cytometry) in which their intended target GPCRs are overexpressed, they should recognize ectodomains of their targeting GPCRs (**Table 1**). We applied each mAb individually to a pre-blocked VirD-GPCR array, measured the corresponding binding signals using fluorescently labeled secondary antibodies, and determined the Z-scores of all the VirD-GPCRs for that mAb on the basis of the standard deviation (S.D.) value for each assay. Using a stringent cut off value of Z-score ≥ 5, only nine of the 20 tested mAbs recognized their intended targets specifically. Two examples of anti-CXCR2 and -CCR7 are shown in **Fig. 2a-b**. Of note, four of them are known to have neutralization activity. One other mAb, anti-ACKR3, not only recognized VirD-ACKR3 as its top 1 target, but also showed Z-scores of 6.5 and 5.9 to VirD-CALCR and VirD-GPR61, respectively, suggesting off-target binding activity. The remainder ten mAbs, however, completely failed to show any detectable binding activities to their intended targets, and some of them bound to many unrelated VirD-GPCRs (i.e., anti-DRD1 in right panel; **Fig. 2a**).

**Table 1.** Summary of the binding specificity tests for 20 commercial mAbs using VirD-GPCR arrays.

| **Symbol** | **Uniport ID** | **Assays on the datasheet** | **VirD array results** | **Z-score** |
| --- | --- | --- | --- | --- |
| CCR7 | P32248 | Neutralization; Flow; IFC | Specific | 16.5 |
| CXCR1 | P25024 | Neutralization; Flow; IHC | Specific | 9.2 |
| CXCR2 | P25025 | Neutralization; Flow; IHC | Specific | 15.6 |
| CXCR5 | P32302 | Neutralization; Flow; IFC; IHC | Specific | 11.4 |
| GPRC5C | Q9NQ84 | Flow | Specific | 16.1 |
| ACKR1 | Q16570 | Flow; IFC | Specific | 18.1 |
| LTB4R | Q15722 | Flow | Specific | 13.4 |
| SSTR2 | P30874 | Flow; IFC; IHC | Specific | 11.8 |
| MRGPRF | Q96AM1 | Flow; WB | Specific | 10.8 |
| ACKR3 | P25106 | Flow; IHC | Specific* | 14.0 |
| DRD1 | P21728 | Flow; IHC | Nonspecific | 2.2 |
| CCR1 | P32246 | Flow | Nonspecific | 1.5 |
| CCR2 | P41597 | Flow, IHC | Nonspecific | -0.5 |
| CCR4 | P51679 | Flow | Nonspecific | 1.4 |
| CCR9 | P51686 | Flow; IFC; IHC | Nonspecific | 0.8 |
| CXCR6 | O00574 | Flow | Nonspecific | -0.7 |
| S1PR1 | P21453 | Flow | Nonspecific | 1.2 |
| APLNR | P35414 | Flow; IHC | Nonspecific | 2.4 |
| P2RY11 | Q96G91 | Flow | Nonspecific | 2.7 |
| P2RY13 | Q9BPV8 | Flow, ELISA, WB | Nonspecific | 1.0 |

Twenty commercial mAbs against 20 GPCR ectodomains were purchased and tested individually on the VirD-GPCR arrays. Z-scores that were calculated for the intended targets of each mAb obtained on the VirD-GPCR arrays are shown. \*VirD-ACKR3 was recognized as the top target with two off-targets, CALCR and GPR61.

**Figure 2.**
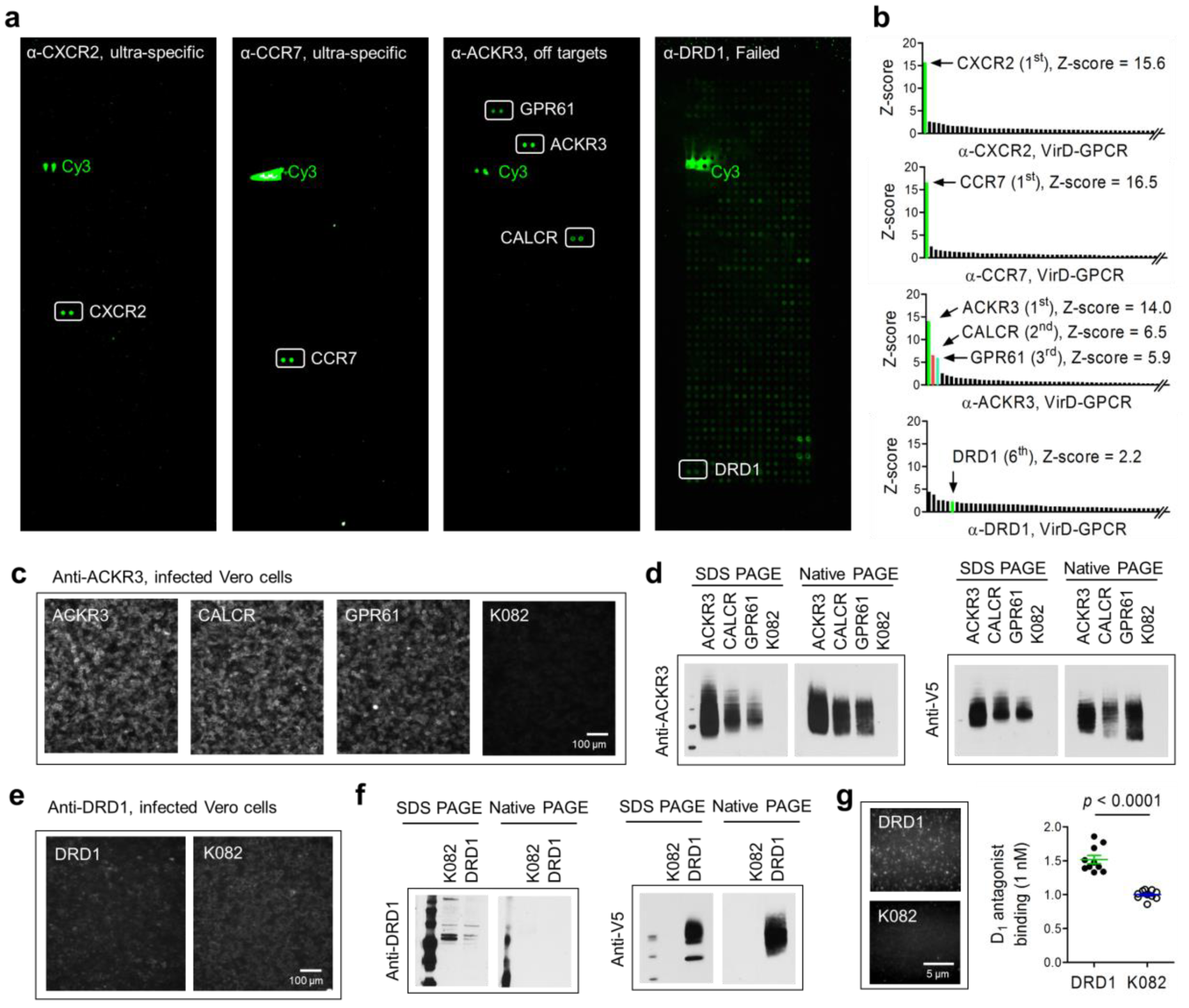
Specificity tests of commercial mAbs on VirD-GPCR arrays. (**a**) Examples of binding signals obtained with commercial mAbs. Anti-CXCR2 and -CCR7 are shown as ultra-specific; anti-ACKR3 can cross-react with CALCR and GPR61; anti-DRD1 completely failed to recognize its target while showing non-specific binding activities to other GPCRs. (**b**) Histograms of Z-scores obtained with three mAbs. Z-scores of the two off- targets identified by anti-ACKR3 are also shown. (**c**) IFA validation of anti-ACKR3 to its off targets in infected Vero cells. K082-infected cells are shown as a negative control. (**d**) Immunoblot analysis also confirmed that anti-ACKR3 can recognize its two off- targets in the cell lysates of infected Vero cells under both denatures and native conditions. (**e-f**) Anti-DRD1 failed to recognize DRD1 in VirD-DRD1-infected cells using IFA (**e**) or IB analyses (**f**). (**g**) Single-molecule imaging using TIRF microscopy to determine interactions between VirD-DRD1 and its canonical ligand D_1_ antagonist. K082 virions were used as a negative control. Quantitative analysis of TIRF imaging demonstrated that VirD-DRD1 showed significantly higher binding signals to D_1_ antagonist than K082 (right panel).

Although an easy explanation is that these mAbs are of poor quality and thus failed in this test, it was also possible that the 11 GPCRs were displayed in the wrong orientation/misfolding in the virions. To explore the latter possibility, we first focused on the off-target binding activity observed with anti-ACKR3 mAb. Using intact Vero cells infected with viruses carrying ACKR3, CALCR, or GPR61, we performed immunofluorescence assays (IFA) with anti-ACKR3. As shown in **Fig. 2c**, the mAb showed strong staining to all three infected cells as compared with the negative control cells infected with K082, a *∆UL27* HSV-1 virus, suggesting that this mAb could recognize CALCR and GPR61 embedded in intact cell membranes. Similarly, this mAb also strongly recognized all three GPCRs in immunoblot (IB) analyses against cell lysates of these infected cells under either native or denatured conditions (**Fig. 2d**). Interestingly, sequence alignment analysis using the ectodomain sequences of the three GPCRs identified a highly conserve 6-mer motif of [NMF]GEL[VTG][RC], suggesting a commonly shared epitope for this mAb. Homology search did not identify any other non-odorant GPCRs that carry a 6-mer peptide with high similarity.

Next, we randomly selected four (i.e., anti-DRD1, -CCR2, -CCR9, and -S1PR1) of the 10 mAbs that completely failed to recognize their intended GPCRs on the VirD-GPCR arrays for the IFA and IB analyses. All of the four mAbs failed these tests; anti-DRD1 is shown in **Fig. 2e-f** as an example (**Fig. 2e-f**; **Supplementary Fig. 2**). The failure of these mAbs was not due to poor expression of their target GPCRs because these GPCRs could be readily detected with anti-V5 (**Fig. 2f**). To determine whether these failed mAbs could recognize linear epitopes, we performed mAb binding assays on HuProt arrays, each comprised of 20,240 human proteins in full-length and denatured using 9 M urea treatment ^18^. Except anti-S1PR1, which was not tested because S1PR1 was not available on HuProt, the rest nine mAbs failed to recognize their intended targets as the top targets. Similarly, none of them recognized their intended targets under native conditions on HuProt arrays (**Supplementary Fig. 3**).

### Functional test of GPCR activity with canonical ligands

To demonstrate further that the ten GPCRs that were not recognized by their respective mAbs were functional/folded correctly, we decided to test their binding activities to their canonical ligands using an imaging approach. Four of the ten ligands were commercially available small molecules with fluorescent labels; the rest were peptide ligands and we labeled them with NHS-conjugated dyes. We immobilized these VirD-GPCRs on a passivated cover slip and incubated with their corresponding ligands at low nanomolar concentrations. Using single-molecule imaging on total internal reflection fluorescence (TIRF) microscopy, we recorded the resulting fluorescent images and compared the binding signals with that of the negative control, K082 virus. Except P2RY13, all the eight VirD-GPCRs that were not recognized by the eight commercial mAbs showed significantly higher binding signals than the K082 control, indicating that they were functional and folded correctly (**Supplementary Fig. 4**). As an example, D_1_ antagonist showed significantly higher binding signals to its canonical receptor DRD1 than K082 (**Fig. 2g**). P2RY13 receptor was not observed to interact with its ligand, ATP, presumably due to the weak affinity in the low micromolar range ^19^. Taken together, the high-content non-odorant VirD-GPCR array was validated as a powerful platform to screen for high quality mAbs against folded GPCRs.

### Specificity test for GPCR ligands

The success of the above approach prompted us to determine whether VirD-GPCR arrays could be used to examine binding specificity of GPCR ligands. As a proof of concept, we chose two peptide ligands, dynorphin A and somatostatin-14 (SRIF-14), to perform specificity tests. As illustrated in **Fig. 3a**, dynorphin A bound to its canonical receptor, OPRD1, as the top receptor with significant signal intensity ^20^. However, SRIF-14 bound to more than 15 VirD-GPCRs with Z-scores ≥ 2 (**Fig. 3b**), including its known receptor, SSTR2 ^21^. To determine whether the observed off-target binding activities of SRIF-14 were not due to an artifact caused by dye-labeling, we employed a cell-based competition assay to examined binding specificity between SRIF-14 and three randomly selected off-target GPCRs, namely GABBR2, NTSR1, and KISS1R, with Z-scores ≥ 2 (**Fig. 3c**). The three VirD-GPCR constructs were used to separately infect Vero cells and VirD-SSTR2 and K082 were also included as positive and negative controls, respectively. The same fluorescently labeled SRIF-14 were added to these infected Vero cells in the absence or presence of cold SRIF-14 (i.e., agonist) or cyclosomatostatin (i.e., antagonist). Quantitative analysis clearly showed that all cell lines infected with the three off-target GPCRs showed significantly higher binding signals than K082-infected cells (upper panel **Fig. 3c**; **Fig. 3d**), comparable to those infected with SSTR2. More importantly, both cold SRIF-14 and cyclosomatostatin (cycloSST) could readily compete off the binding signals of Cy5-labeled SRIF-14 on the cells infected with the three off-target GPCRs, suggesting ligand-specific interactions (middle and lower panels; **Fig. 3c**; **Fig. 3d**).

**Figure 3.**
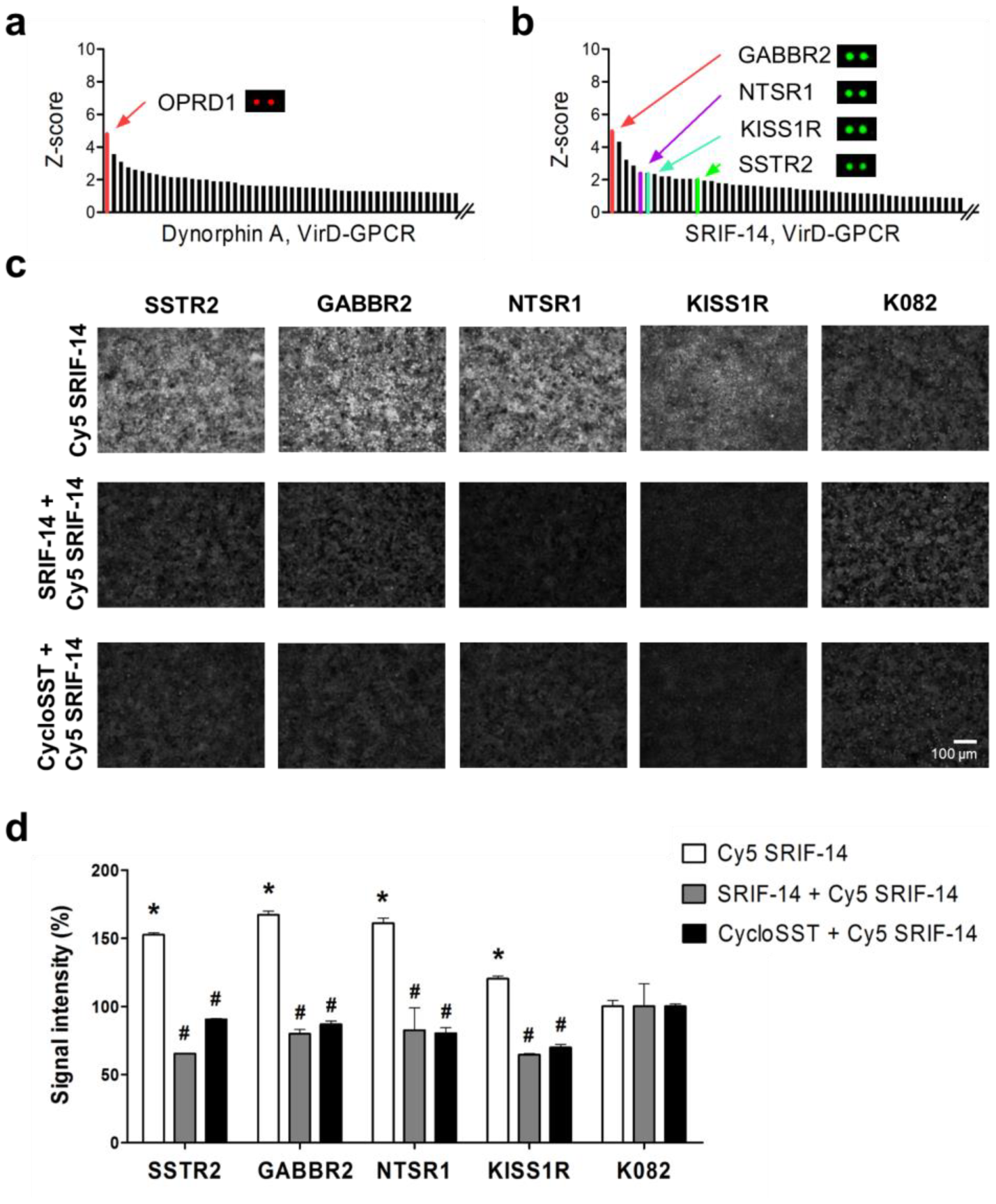
Identification and cell-based validation of peptide ligand-GPCR interactions. (**a**) Commercially available Dynorphin A was Cy5-labeled and probed to a VirD-GPCR array. Quantitative analysis showed that it bound to VirD-OPRD1 with the highest Z-score. (**b**) A commercially available peptide SRIF-14 was Cy3-labeled and probed to a VirD-GPCR array. Quantitative analysis revealed that it bound to several unexpected off-targets in addition to its canonical receptor, SSTR2. (**c**) Vero cells were separately infected with SSTR2, GABBR2, NTSR1, KISS1R, or K082 virus. Infected cells were then incubated with Cy5-labeled SRIF-14 at 8 μM in the absence (upper panel) or presence of cold SRIF-14 (middle panel) or cyclosomatostatin (cycloSST, lower panel). (**d**) Quantitative analysis of binding signals. Each binding assay was performed in triplicate and the obtained binding signals were normalized to those of the K082 controls. * *P* < 0.05, comparison between VirD-GPCRs and K082 in the absence of competitor ligands; # *P* < 0.001, comparison between binding signals obtained in the absence and presence of the competitor ligands.

Because the array- or cell-based assays can only offer an end-point measurement, we decided to obtain true binding kinetics using a previously reported virion-oscillator approach ^22, 23^. It is a label-free plasmonic imaging technique that can quantify the ligand binding-induced mobility change of the virions anchored on the surface of the sensor chip via a flexible molecular linker and reveal the binding kinetics of small molecule ligands. VirD-GABBR2, -NTSR1, and -KISS1R were separately attached to a gold-coated cover slip via a flexible polyethylene glycol (PEG) linker (**Fig. 4a**). Again, VirD-SSTR2 and K082 were included as the positive and negative controls, respectively. Using surface plasmon resonance (SPR) to monitor voltage-induced oscillation of the VirD-GPCRs, we recorded oscillation amplitude changes of VirD-GPCRs as a result of association and dissociation of SRIF-14 in a real-time, label-free fashion. As illustrated in **Fig. 4b-f**, all four tested VirD-GPCRs showed typical association and dissociation curves in a dose-dependent manner, which allowed us to determine the corresponding *k*_*a*_ and *k*_*d*_ values (**Supplemental Table 1**). As expected, K082 did not show any detectable binding kinetics even in the presence of 8 μM SIRF-14 (**Fig. 4f**). Binding affinity *K_D_* values were thus calculated for SSTR2, GABBR2, NTSR1, and KISS1R to be 11.2 nM, 0.4 μM, 2.6 μM, and 25.0 μM, respectively. These independent experiments confirmed the off-target binding activities of SIRF-14 observed on VirD-GPCR arrays.

**Figure 4.**
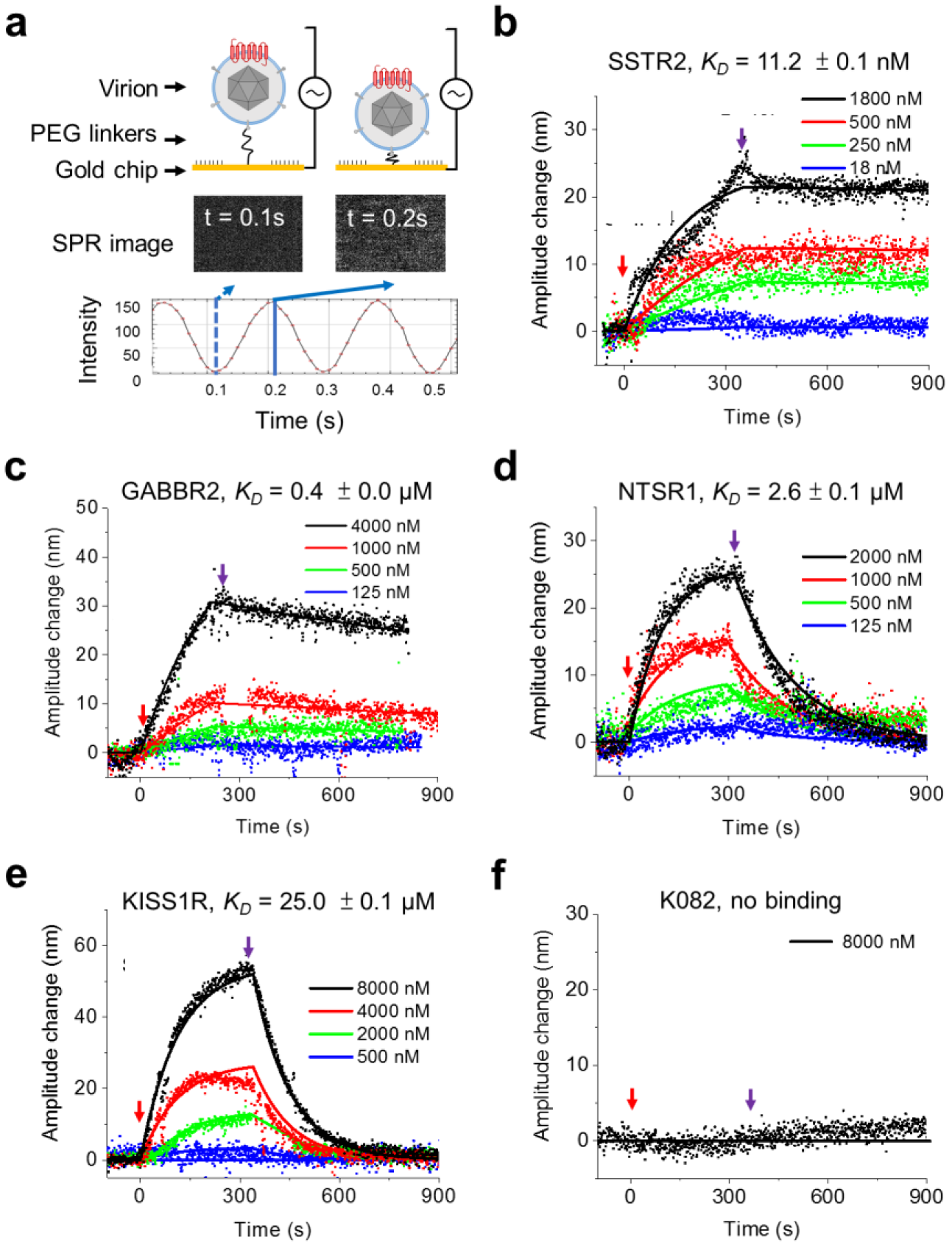
Binding kinetics of SRIF-14 to SSTR2, GABBR2, NTSR1, KISS1R, and K082 virions. (**a**) Principle of binding kinetics measurement using virion-oscillator device. (**b-f**) Binding kinetics of SRIF-14 to its canonical receptor, SSTR2, and three newly discovered off-target GPCRs. The red and purple arrows mark the starting time points of the association and the dissociation phases, respectively. Binding curves were fit using the first order kinetics model (solid lines). The calculated affinity values range from 11.2 nM (SSTR2) to 25.0 μM (KISS1R), while K082 virion showed no detectable binding activity to the SRIF-14. Detailed *k*_*a*_, *k*_*d*_, and *K*_*D*_ values were listed in **Supplementary table 1**.

### Discovery and characterization of GPCR targets for group B Streptococcus

In recent years, several elegant studies reported that both Gram-negative and -positive bacterial pathogens, such as *Neisseria meningitidis* and *Streptococcus pneumoniae*, could utilize human GPCRs (e.g., ADRB2 and PTAFR) as receptors to penetrate human epithelial cells ^24, 25^. *Streptococcus agalactiae* (a.k.a. group B strep or GBS) is the most common Gram-positive organism causing neonatal meningitis by penetrating human blood brain barriers (BBB). However, its host receptor has remained elusive. To explore the possibility that GBS may exploit host GPCRs as a means to penetrate BBB, we probed the VirD-GPCR array with fluorescently labeled live GBS K79 (a strain isolated from a neonate with meningitis) in duplicate to discover potential host receptors. As a comparison, a Gram-positive non-pathogenic bacterium, *Streptococcus gordonii*, which does not penetrate BBB, was used as a negative control. Five VirD-GPCRs, namely GPR101, GPR148, LHCGR, CysLTR1, and LGR5, showed significantly higher binding signals to K79 than *S. gordonii* in a reproducible manner (**Fig. 5a**).

**Figure 5.**
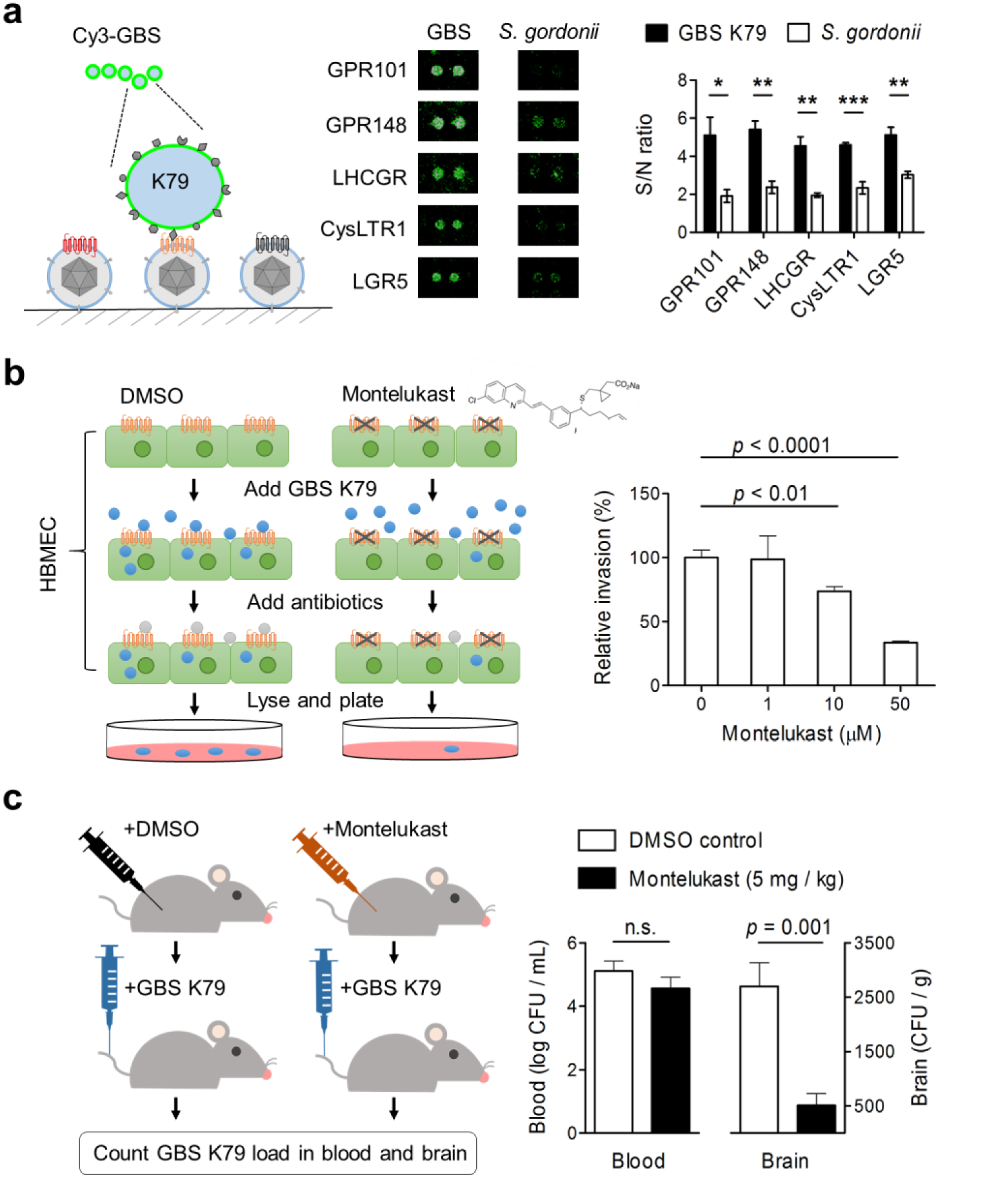
Discovery and validation of a new GPCR receptor for GBS. (**a**) Five VirD-GPCRs were identified as potential candidate receptors. A clinical strain K79 of GBS was Cy3-labeled and probed to a VirD-GPCR array. In parallel, *S. gondonii* was used as a non-pathogenic negative control (middle). Quantitative analysis of the binding signals from the two bacteria identified five GBS-specific GPCRs (right). (**b**) *In vitro* validation of CysLTR1. HBMEC cells were pretreated with Montelukast to block CysLTR1, followed by incubation with GBS. After washes, antibiotics were added to kill free bacteria and HBMEC cells were lysed and plated onto blood agar plates. After overnight incubation, numbers of GBS colonies were counted. As compared with the DMSO-treated negative control, Montelukast showed a dose-dependent inhibition of GBS invasion into HBMEC cells. (**c**) *In vivo* validation of CysLTR1. A group of mice were each intraperitoneally administered Montelukast (n = 6) or DMSO (n = 5). After 2 hr each mouse received 1×10^8^ CFU of GBS (K79) via the tail vein injection. One hr later, blood and homogenized brains were collected and plated for bacterial counts. Using the same colony formation method, administration of Montelukast reduced GBS brain infection in the mice by an average of 81% as compared to the DMSO controls (right). No significant differences in the levels of GBS counts were observed in the blood between the two groups.

Of the five identified potential GBS receptors, GPR101, GPR148 and LGR5 are “orphan” receptors without identified canonical ligands. LHCGR is mainly expressed in ovary and testis and binds luteinizing hormone; mutations in this gene are known to cause infertility ^26^. Therefore, these four candidates are either difficult to pursue or less relevant to the pathogenesis of meningitis. The last candidate receptor, CysLTR1, recognizes cysteinyl leukotrienes and is an attractive candidate for several reasons. First, its activation is associated with increased permeability of BBB, as well as promotion of the movement of leukocytes from the blood into brain tissues in animal models ^27^. Second, its activation may also increase the entry of leukocyte-borne viruses, such as HIV-1, into brain tissue ^28^. Third, a well-established antagonist, Montelukast, specifically inhibits CysLTR1 but not its homolog CysLTR2 ^29^. Importantly, recent studies have demonstrated that Montelukast could protect against hippocampus injury induced by transient ischemia and reperfusion in rats ^30^.

To validate further the discovery of CysLTR1 as a potential target of GBS, we first employed cultured human brain microvascular endothelial cells (HBMEC) cells to evaluate the role of endogenous CysLTR1 in GBS penetration. HBMEC is the major component of the blood-brain barrier and CysLTR1 is known to be expressed in this cell line ^31^. Monolayers of HBMEC were pre-treated with Montelukast at different concentrations for 1 hr, washed, and infected with GBS K79 (see Methods for more details). DMSO-treated cells were used as a vehicle control. After removal of unbound bacteria with several washes, antibiotics (gentamicin and penicillin) were added to the cells to kill extracellular bacteria. To evaluate GBS penetration to the cells under different conditions, cells were lysed, diluted, and plated onto blood agar plates. Colonies formed on the agar plates were counted and used as a proxy to evaluate GBS penetration of the blood-brain barrier (see Methods). After normalization to the DMSO controls, it was clear that Montelukast significantly inhibited GBS penetration to HBMEC cells in a dose-dependent fashion. In fact, as much as 77% GBS penetration was inhibited by 50 μM Montelukast (**Fig. 5b**). These results corroborated our VirD-GPCR array results that CysLTR1 might play an important role in GBS penetration of the blood-brain barrier.

To further demonstrate the role of CysLTR1 in GBS penetration into the brain *in vivo*, we employed a mouse model of experimental hematogenous meningitis (**Fig. 5c**). A group of six mice were each intraperitoneally administered Montelukast (5 mg/kg), while another group of five mice received DMSO as a vehicle control. After 2 hr each mouse received 1×10^8^ CFU of GBS (K79) via the tail vein injection. One hour later, blood was collected and plated for bacteria counts. Immediately following the blood collection, mice were transcardially perfused to remove the remaining body blood and the brains were removed, weighed, homogenized, and plated for bacterial counts (see Methods for more details). We found that the administration of Montelukast reduced the GBS infection in the mouse brains by an average of 81% as compared to that of the DMSO controls. It is important to note that no significant differences of GBS counts were observed in the blood between the two groups, indicating that decreased GBS penetration into the brain was not the result of having less bacterial counts in the blood at the time of collecting the brain specimens (**Fig. 5c**). Taken together, these experiments demonstrated that using the VirD-GPCR array as an unbiased screening platform, we successfully identified CysLTR1 as a receptor for GBS penetration into host cells both *in vitro* and *in vivo*.

## DISCUSSION

The Virion Display (VirD) method was developed to display membrane protein on the lipid envelope of HSV-1 virion. To take a full advantage of VirD technology, we built a high-content non-odorant GPCR array, comprised of 315 human GPCRs. This is the first array of its kind, covering >85% of all annotated non-odorant GPCRs, which are preferred drug targets. Using this high-content VirD-GPCR array as a tool, we were able to characterize the specificity of commercial antibodies and canonical ligands, and to identify new receptors involved in host-pathogen interactions.

Unlike the existing display systems, such as liposomes, nanodiscs ^32, 33^, and cell membrane fractions ^7, 8^, GPCRs displayed in the VirD-GPCR system are most likely to orientate uniformly with preserved conformation, as demonstrated in this study and Hu et al ^9^. In addition, copies of the GPCR molecules displayed on a single virion are more uniform because they are expressed as viral proteins and assembled by the virus machinery. Another display technology generates virus-like particles (VLPs) to maintain native conformation of membrane proteins. Unlike VirD, however, VLPs do not carry any genetic information and are not self-renewable. Therefore, transient transfection is needed every time to regenerate VLPs, raising concerns of reproducibility ^34^. In addition, VirD-GPCRs are much more stable, because they are carried by HSV-1 viral particles, and are more amenable to large-scale production.

Another unique advantage of VirD technology that has not yet been fully exploited is the flexibility in choice of cell lines for VirD production, since HSV-1 can infect many cell line *in vitro* ^35^. For example, we have used HEL, Vero, HeLa, and HEK293T cells for virion production to obtain the optimal GPCR expression (**Supplementary Fig. 1 a-b**). This can be very important for GPCRs that require tissue-specific posttranslational modifications (e.g., glycosylation) to maintain their native functions. Moreover, the infectious stocks of the VirD-GPCR collection generated in this study can be readily used to infect many primate cell lines, facilitating downstream mechanistic studies, such as calcium imaging and β-arrestin recruitment. In its current form, a single GPCR is displayed per virion. However, many GPCRs are known to form obligate heterodimers (e.g., GABA receptor GABBR1/GABBR2) or even multimers to achieve additional regulation ^36, 37^. Because of the flexibility of the VirD platform, we can co-infect human cells with more than one type of VirD-GPCR to produce chimeric heterodimer or multimer VirD-GPCRs.

Monoclonal antibodies (mAbs) are fast growing therapeutic modalities and account for ~30% of the new drugs approved by the FDA in recent years ^38, 39^. Indeed, FDA has just approved the first mAb-based drug targeting CGRPR ^11^. However, mAb specificity has been one of the major concerns and antibodies of poor performance have wasted about half of the spending on the protein-binding reagents ^15–17^. This is particularly true for the GPCR community because a major hurdle to generating antibodies against GPCRs is obtaining folded GPCRs as antigens for animal immunization or display platform-based discovery. Since the conformation of VirD-GPCRs is preserved, VirD-GPCR arrays offer a high-throughput platform to screen for functional antibodies of high specificity with therapeutic potentials. Indeed, we purposefully selected 20 commercial mAbs that were claimed by the manufactures to recognize the ectodomains of 20 different human GPCRs. However, only nine mAbs turned out to be ultra-specific, four of which are known to have neutralization activities, indicating that they recognize functional epitopes on VirD-GPCR arrays. Although 10 commercial mAbs failed to recognize their intended targets in VirD-GPCR binding assays, nine of the 10 VirD-GPCRs could bind their canonical ligands as demonstrated with TIRF microscopy, indicating that these GPCRs are all folded correctly. Therefore, we believe that VirD-GPCR array is an ideal platform to identify the specificity and potential therapeutic value of mAbs against native and fully folded GPCRs.

Another application of the VirD-GPCR array is to profile the specificity of ligands. Cy5-labeled Dynorphin A bound to one of its canonical receptors on the VirD-GPCR array with high specificity, but Cy3-labeled SRIF-14 bound to multiple unanticipated receptors (e.g., GABBR2, NTSR1, and KISS1R) in addition to its canonical receptor, SSTR2. These off-target binding activity was further validated using the virion-oscillator technology, even though the binding affinity of SRIF-14 to the off-target GPCRs was in the low μM to high nM range. Peptide ligands of low affinity are a common property of many GPCRs, such as GPR55 binds to cannabinoid ^40^. However, detection of binding activity is only the first step to identify a new ligand for GPCRs; *in vitro* and *in vivo* functional assays, such as calcium imaging, PRESTO-Tango assays, and animal-based studies, will be required to fully characterize these new interactions. Nevertheless, VirD-GPCR arrays can offer a rapid, comprehensive survey for specificity of ligand-GPCR interactions.

Although GPCRs are mostly known for their involvement in a wide variety of physiological processes, such as inflammation, neurotransmission, and autonomic nervous system transmission, recent studies also demonstrated that some GPCRs play an important role in pathogen-host interactions ^24, 25^. GPCRs can serve as an entry point for pathogens because GPCRs undergo an internalization process via β-arrestins following activation. To explore the possibility of using the VirD-GPCR array to identify novel receptors for bacterial pathogens, we probed labeled GBS on a VirD-GPCR array and readily identified several GPCR candidates that showed specific interactions as compared with a negative control bacterium. By focusing on one candidate receptor, CysLTR1, we demonstrated that this newly identified GPCR indeed played a crucial role in GBS invasion of BBB both *in vitro* and *in vivo*. We believe that this approach can be easily applied to many other microbial pathogens to better understand the molecular mechanisms of pathogen invasion and provide novel therapeutic targets.

## METHODS

Methods, including statements of data availability and any associated accession codes and references, are available in the online version of the paper.

## ACKNOWLEDGEMENTS

This work was supported in part by the NIH / NCI IMAT R33-CA186790-01A1 (H.Z. and P.J.D.), NIH TCNP Roadmap (H.Z.), NIH NS091102, AI84984, AI113273 and AI126176 (K.S.K), NIH R01NS054791 (X.D.), Johns Hopkins University Discovery Awards (J.X.), Hamilton Innovation Award (J.X.), Taiwanese Government Scholarship Award (SC.W.), CDI Laboratories, and MOST Postdoctoral Fellowship -Taiwan (G.S.). We would like to thank the following colleagues for their assistance and support: David Leib (Dartmouth University) for the gift of KOS-37 BAC; David Johnson at Oregon Health and Science University for the generous gift of anti-gD antibody, DL6. We also thank Jen Tullman and Wade Gibson (Johns Hopkins University) for helpful discussions on virion purification and BAC recombineering method.

## AUTHOR CONTRIBUTIONS

G.S., SC.W., G.M., S.L., A.P., D.P., B.H., D.G., P.R., P.J.D., and L.W. performed experimental work. G.S., H.Z., P.J.D., K.S.K., N.T., S.W., D.E., I.P., X.D., J.X., SC.W., G.M., and A.P. contributed to manuscript preparation. N.T., S.W., D.E., I.P., X.D., and J.X. contributed their expertise and supervision to the work. H.Z., P.J.D., and K.S.K. conceived the idea and supervised the entire project. H.Z. and G.S. wrote the manuscript.

## COMPETING INTERESTS

H.Z., D.E., and I.P. are cofounders and shareholders of CDI Laboratories Inc. P.R. is an employee of CDI Laboratories Inc. H.Z., G.S. and P.J.D. are consultants to CDI Laboratories Inc.

## MATERIALS & CORRESPONDENCE

H.Z., P.J.D., and K.S.K.

## REFERENCES

1. Hauser, A.S., Attwood, M.M., Rask-Andersen, M., Schioth, H.B. & Gloriam, D.E. Trends in GPCR drug discovery: new agents, targets and indications. Nature reviews. Drug discovery 16, 829–842 (2017).

2. Hauser, A.S. et al. Pharmacogenomics of GPCR Drug Targets. Cell 172, 41–54 e19 (2018).

3. Sriram, K. & Insel, P.A. G Protein-Coupled Receptors as Targets for Approved Drugs: How Many Targets and How Many Drugs? Molecular pharmacology 93, 251–258 (2018).

4. Duvernay, M.T., Filipeanu, C.M. & Wu, G. The regulatory mechanisms of export trafficking of G protein-coupled receptors. Cellular signalling 17, 1457–1465 (2005).

5. Jacoby, E., Bouhelal, R., Gerspacher, M. & Seuwen, K. The 7 TM G-protein-coupled receptor target family. ChemMedChem 1, 761–782 (2006).

6. Milic, D. & Veprintsev, D.B. Large-scale production and protein engineering of G protein-coupled receptors for structural studies. Frontiers in pharmacology 6, 66 (2015).

7. Fang, Y., Peng, J., Ferrie, A.M. & Burkhalter, R.S.. Air-stable G protein-coupled receptor microarrays and ligand binding characteristics. Analytical chemistry 78, 149–155 (2006).

8. Fang, Y., Frutos, A.G. & Lahiri, J. Membrane protein microarrays. Journal of the American Chemical Society 124, 2394–2395 (2002).

9. Hu, S. et al. VirD: a virion display array for profiling functional membrane proteins. Analytical chemistry 85, 8046–8054 (2013).

10. Gierasch, W.W. et al. Construction and characterization of bacterial artificial chromosomes containing HSV-1 strains 17 and KOS. Journal of virological methods 135, 197–206 (2006).

11. Goadsby, P.J. et al. A Controlled Trial of Erenumab for Episodic Migraine. The New England journal of medicine 377, 2123–2132 (2017).

12. Weller, M.G. Quality Issues of Research Antibodies. Analytical chemistry insights 11, 21–27 (2016).

13. Bordeaux, J. et al. Antibody validation. BioTechniques 48, 197–209 (2010).

14. Schonbrunn, A. Editorial: Antibody can get it right: confronting problems of antibody specificity and irreproducibility. Mol Endocrinol 28, 1403–1407 (2014).

15. Baker, M. Antibody anarchy: A call to order. Nature 527, 545–551 (2015).

16. Bradbury, A. & Pluckthun, A. Reproducibility: Standardize antibodies used in research. Nature 518, 27–29 (2015).

17. Herrera, M., Sparks, M.A., Alfonso-Pecchio, A.R., Harrison-Bernard, L.M. & Coffman, T.M. Lack of specificity of commercial antibodies leads to misidentification of angiotensin type 1 receptor protein. Hypertension 61, 253–258 (2013).

18. Venkataraman, A. et al. A toolbox of immunoprecipitation-grade monoclonal antibodies to human transcription factors. Nature methods (2018).

19. Marteau, F. et al. Pharmacological characterization of the human P2Y13 receptor. Molecular pharmacology 64, 104–112 (2003).

20. Zhang, S. et al. Dynorphin A as a potential endogenous ligand for four members of the opioid receptor gene family. The Journal of pharmacology and experimental therapeutics 286, 136–141 (1998).

21. Liapakis, G. et al. Identification of ligand binding determinants in the somatostatin receptor subtypes 1 and 2. The Journal of biological chemistry 271, 20331–20339 (1996).

22. Fang, Y., Chen, S., Wang, W., Shan, X. & Tao, N. Real-time monitoring of phosphorylation kinetics with self-assembled nano-oscillators. Angew Chem Int Ed Engl 54, 2538–2542 (2015).

23. Shan, X. et al. Detection of charges and molecules with self-assembled nano-oscillators. Nano letters 14, 4151–4157 (2014).

24. Coureuil, M. et al. Meningococcus Hijacks a beta2-adrenoceptor/beta-Arrestin pathway to cross brain microvasculature endothelium. Cell 143, 1149–1160 (2010).

25. Radin, J.N. et al. beta-Arrestin 1 participates in platelet-activating factor receptor-mediated endocytosis of Streptococcus pneumoniae. Infection and immunity 73, 7827–7835 (2005).

26. Wu, S.M. & Chan, W.Y. Male pseudohermaphroditism due to inactivating luteinizing hormone receptor mutations. Archives of medical research 30, 495–500 (1999).

27. Lenz, Q.F. et al. Cysteinyl leukotriene receptor (CysLT) antagonists decrease pentylenetetrazol-induced seizures and blood-brain barrier dysfunction. Neuroscience 277, 859–871 (2014).

28. Bertin, J., Jalaguier, P., Barat, C., Roy, M.A. & Tremblay, M.J. Exposure of human astrocytes to leukotriene C4 promotes a CX3CL1/fractalkine-mediated transmigration of HIV-1-infected CD4(+) T cells across an in vitro blood-brain barrier model. Virology 454–455, 128–138 (2014).

29. Nothacker, H.P. et al. Molecular cloning and characterization of a second human cysteinyl leukotriene receptor: discovery of a subtype selective agonist. Molecular pharmacology 58, 1601–1608 (2000).

30. Saad, M.A., Abdelsalam, R.M., Kenawy, S.A. & Attia, A.S. Montelukast, a cysteinyl leukotriene receptor-1 antagonist protects against hippocampal injury induced by transient global cerebral ischemia and reperfusion in rats. Neurochemical research 40, 139–150 (2015).

31. Zhu, L., Maruvada, R., Sapirstein, A., Peters-Golden, M. & Kim, K.S. Cysteinyl leukotrienes as novel host factors facilitating Cryptococcus neoformans penetration into the brain. Cellular microbiology 19 (2017).

32. Shen, H.H., Lithgow, T. & Martin, L. Reconstitution of membrane proteins into model membranes: seeking better ways to retain protein activities. International journal of molecular sciences 14, 1589–1607 (2013).

33. Borch, J. & Hamann, T. The nanodisc: a novel tool for membrane protein studies. Biological chemistry 390, 805–814 (2009).

34. Willis, S. et al. Virus-like particles as quantitative probes of membrane protein interactions. Biochemistry 47, 6988–6990 (2008).

35. Karasneh, G.A. & Shukla, D. Herpes simplex virus infects most cell types in vitro: clues to its success. Virology journal 8, 481 (2011).

36. Vischer, H.F., Castro, M. & Pin, J.P. G Protein-Coupled Receptor Multimers: A Question Still Open Despite the Use of Novel Approaches. Molecular pharmacology 88, 561–571 (2015).

37. Gurevich, V.V. & Gurevich, E.V. How and why do GPCRs dimerize? Trends in pharmacological sciences 29, 234–240 (2008).

38. Mullard, A. 2017 FDA drug approvals. Nature reviews. Drug discovery 17, 81–85 (2018).

39. Mullard, A. 2016 FDA drug approvals. Nature reviews. Drug discovery 16, 73–76 (2017).

40. Kapur, A. et al. Atypical responsiveness of the orphan receptor GPR55 to cannabinoid ligands. The Journal of biological chemistry 284, 29817–29827 (2009).

